# A widely distributed hydrogenase oxidises atmospheric H_2_ during bacterial growth

**DOI:** 10.1101/2020.04.14.040717

**Authors:** Zahra F. Islam, Caitlin Welsh, Katherine Bayly, Rhys Grinter, Gordon Southam, Emma J. Gagen, Chris Greening

**Author notes:** Correspondence can be addressed to: Assoc Prof Chris Greening.

## Abstract

Diverse aerobic bacteria persist by consuming atmospheric hydrogen (H_2_) using group 1h [NiFe]-hydrogenases. However, other hydrogenase classes are also distributed in aerobes, including the group 2a [NiFe]-hydrogenase. Based on studies focused on Cyanobacteria, the reported physiological role of the group 2a [NiFe]-hydrogenase is to recycle H_2_ produced by nitrogenase. However, given this hydrogenase is also present in various heterotrophs and lithoautotrophs lacking nitrogenases, it may play a wider role in bacterial metabolism. Here we investigated the role of this enzyme in three species from different phylogenetic lineages and ecological niches: *Acidithiobacillus ferrooxidans* (phylum Proteobacteria), *Chloroflexus aggregans* (phylum Chloroflexota), and *Gemmatimonas aurantiaca* (phylum Gemmatimonadota). qRT-PCR analysis revealed that the group 2a [NiFe]-hydrogenase of all three species is significantly upregulated during exponential growth compared to stationary phase, in contrast to the profile of the persistence-linked group 1h [NiFe]-hydrogenase. Whole-cell biochemical assays confirmed that all three strains aerobically respire H_2_ to sub-atmospheric levels, and oxidation rates were much higher during growth. Moreover, the oxidation of H_2_ supported mixotrophic growth of the carbon-fixing strains *C. aggregans* and *A. ferrooxidans.* Finally, we used phylogenomic analyses to show that this hydrogenase is widely distributed and is encoded by 13 bacterial phyla. These findings challenge the current persistence-centric model of the physiological role of atmospheric H_2_ oxidation and extends this process to two more phyla, Proteobacteria and Gemmatimonadota. In turn, these findings have broader relevance for understanding how bacteria conserve energy in different environments and control the biogeochemical cycling of atmospheric trace gases.

## Introduction

Aerobic bacteria mediate the biogeochemically and ecologically important process of atmospheric hydrogen (H_2_) oxidation [1]. Terrestrial bacteria constitute the largest sink of this gas and mediate the net consumption of approximately 70 million tonnes of atmospheric H_2_ per year [2, 3]. The energy derived from this process appears to be critical for sustaining the productivity and biodiversity of ecosystems with low organic carbon inputs [4–9]. Atmospheric H_2_ oxidation is thought to be primarily mediated by group 1h [NiFe]-hydrogenases, a specialised oxygen-tolerant, high-affinity class of hydrogenases [4, 10–13]. To date, aerobic heterotrophic bacteria from four distinct bacterial phyla, the Actinobacteriota [10, 12, 14, 15], Acidobacteriota [16, 17], Chloroflexota [18], and Verrucomicrobiota [19], have been experimentally shown to consume atmospheric H_2_ using this enzyme. This process has been primarily linked to energy conservation during persistence. Reflecting this, the expression and activity of the group 1h hydrogenase is induced by carbon starvation across a wide range of species [10, 12, 18, 20–23]. Moreover, genetic deletion of hydrogenase structural genes results in impaired long-term survival of *Mycobacterium smegmatis* cells and *Streptomyces avermitilis* spores [20, 21, 24, 25].

Genomic and metagenomic surveys have suggested that other uptake hydrogenases are widely distributed among aerobic bacteria and potentially have a role in atmospheric H_2_ uptake [4, 26]. These include the widely distributed group 2a [NiFe]-hydrogenases. This hydrogenase class has primarily been investigated in Cyanobacteria, where it is encoded by most diazotrophic strains; the enzyme recycles H_2_ released as a by-product of the nitrogenase reaction and inputs the derived electrons into the respiratory chain [27–30]. However, according to HydDB, group 2a hydrogenases are also encoded by isolates from at least eight other phyla [26], spanning both obligate organoheterotrophs (e.g. *Mycobacterium*, *Runella*, *Gemmatimonas*) and obligate lithoautotrophs (e.g. *Acidithiobacillus*, *Nitrospira*, *Hydrogenobacter*) [12, 31, 32]. In *M. smegmatis*, this enzyme has a sufficiently high apparent affinity to oxidise H_2_ even at sub-atmospheric levels [12, 23] and is maximally expressed during transitions between growth and persistence [23, 33]. In common with the group 1h hydrogenase also encoded by this bacterium, the group 2a hydrogenase requires potential electron-relaying iron-sulfur proteins for activity [34] and is obligately linked to the aerobic respiratory chain [23]. However, it remains unclear if atmospheric H_2_ oxidation by the group 2a hydrogenase reflects a general feature of the enzyme or instead is a specific adaptation of the mycobacterial respiratory chain.

In this study, we investigated whether group 2a [NiFe]-hydrogenases play a general role in atmospheric H_2_ consumption. To do so, we studied this enzyme in three species, *Gemmatimonas aurantiaca*, *Acidithiobacillus ferrooxidans*, and *Chloroflexus aggregans*, that differ in their phylogenetic affiliation, ecological niches, and metabolic strategies. The obligate chemoorganoheterotroph *G. aurantiaca* (phylum Gemmatimonadota) was originally isolated from a wastewater treatment plant and to date has not been shown to utilise H_2_ [35, 36]. The obligate chemolithoautotroph *A. ferrooxidans* (phylum Proteobacteria) was originally isolated from acidic coal mine effluent, and has been extensively studied for its energetic flexibility, including the ability to grow exclusively on H_2_ [32, 37, 38]. The metabolically flexible *C. aggregans* (phylum Chloroflexota), a facultative chemolithoautotroph and anoxygenic photoheterotroph, was originally isolated from a Japanese hot spring and is capable of hydrogenotrophic growth [39–41]. The organisms differ in their carbon dioxide fixation pathways, with *A. ferrooxidans* mediating the Calvin-Benson cycle *via* two RuBisCO enzymes, *C. aggregans* encoding the 3-hydroxypropionate cycle [38, 42, 43], and *G. aurantiaca* unable to fix carbon dioxide [35]. While all three species have previously been shown to encode group 2a [NiFe]-hydrogenases [4, 38], it is unknown whether they can oxidise atmospheric H_2_ oxidation. To resolve this, we investigated the expression, activity, and role of this enzyme in axenic cultures of the three species.

## Materials and Methods

### Bacterial growth conditions

*Gemmatimonas aurantiaca* (DSM 14586), *Acidithiobacillus ferrooxidans* (DSM 14882), and *Chloroflexus aggregans* (DSM 9486) were imported from DSMZ. All cultures were routinely aerobically maintained in 120 mL glass serum vials with treated lab-grade butyl rubber stoppers, unless otherwise stated. Broth cultures of *G. aurantiaca* were grown in 30 mL of NM1 media as previously described [44] and incubated at 30°C at an agitation speed of 180 rpm in a New Brunswick Scientific Excella E24 incubator. Cultures of *C. aggregans* were maintained chemoheterotrophically in 30 mL of 1/5 PE media, as previously described [39], and incubated at 55°C at an agitation speed of 150 rpm in an Eppendorf 40 Incubator in the dark. Cultures of *A. ferrooxidans* were maintained in 30 mL DSMZ medium 882 supplemented with an additional 13 g L^−1^ of FeSO_4_.7H_2_O (pH 1.2) and incubated at 30°C at an agitation speed of 180 rpm in a New Brunswick Scientific Excella E24 incubator. To assess whether bacterial growth was enhanced by the presence of H_2_ for each species, ambient air headspaces were amended with either 1% or 10% H_2_ (*via* 99.999% pure H_2_ gas cylinder). Growth was monitored by determining the optical density (OD_600_) of periodically sampled 1 mL extracts using an Eppendorf BioSpectrophotometer.

### RNA extraction

Triplicate 30 mL cultures of *G. aurantiaca*, *A. ferrooxidans* and *C. aggregans* were grown synchronously in 120 mL sealed serum vials. Whereas one set of triplicate cultures were grown in an ambient air headspace, another set was grown in an ambient air headspace supplemented with H_2_ to a final concentration of 10% v/v (*via* a 99.999% pure H_2_ cylinder). Cultures were grown to either exponential phase (OD_600_ 0.05 for *G. aurantiaca*; OD_600_ 0.1 for *C. aggregans*; OD_600_ 0.05 for *A. ferrooxidans*) or stationary phase (Day 10 for *G. aurantiaca*; Day 4 for *C. aggregans*; Day 14 for *A. ferrooxidans*). For *G. aurantiaca and C. aggregans*, cells were then quenched using a glycerol-saline solution (−20°C, 3:2 v/v), harvested by centrifugation (20,000 × *g*, 30 min, −9°C), resuspended in 1 mL cold 1:1 glycerol:saline solution (−20°C), and further centrifuged (20,000 × *g*, 30 min, −9°C). Briefly, resultant cell pellets were resuspended in 1 mL TRIzol Reagent (Thermo Fisher Scientific), mixed with 0.1 mm zircon beads (0.3 g), and subject to beat-beating (five cycles, 4000 rpm, 30 s) in a Mini-Beadbeater 96 (Biospec) prior to centrifugation (12,000 × *g*, 10 min, 4°C). Total RNA was extracted using the phenol-chloroform method as per manufacturer’s instructions (TRIzol Reagent User Guide, Thermo Fisher Scientific) and resuspended in diethylpyrocarbonate (DEPC)-treated water. RNA was treated using the TURBO DNA-free kit (Thermo Fisher Scientific) as per manufacturer’s instructions. RNA from *A. ferrooxidans* was extracted using a previously described extraction method optimised for acid mine drainage microorganisms [45]. RNA concentration and purity were confirmed using a NanoDrop ND-1000 spectrophotometer.

### Quantitative RT-PCR

Quantitative reverse transcription PCR (qRT-PCR) was used to determine the expression profile of all hydrogenase genes present in each species during different growth phases with and without supplemental H_2_. cDNA was synthesised using a SuperScript III First-Strand Synthesis System kit for qRT-PCR (Thermo Fisher Scientific) with random hexamer primers, as per manufacturer’s instructions. For all three species, the catalytic subunit gene of the group 2a [NiFe]-hydrogenase (*hucL*) was targeted. In addition, the catalytic subunits of the additional [NiFe]-hydrogenases of *C. aggregans* (group 3d, *hoxH*) and *A. ferrooxidans* (group 1e, *hyiB*; group 3b, *hyhL*) were also targeted. Quantitative RT-PCR was performed using a LightCycler 480 SYBR Green I Master Mix (Roche) as per manufacturer’s instructions in 96-well plates and conducted in a LightCycler 480 Instrument II (Roche). Primers used in the study **(Table S1)** were designed using Primer3 [46]. Hydrogenase expression data was normalised to housekeeping genes for each species (16S rRNA gene for *G. aurantiaca* and *C. aggregans*; DNA-directed RNA polymerase subunit beta gene *rpoC* for *A. ferrooxidans*). Threshold cycle values (C_T_) were normalised to the expression of the housekeeping gene in exponential phase under ambient air conditions. All biological triplicate samples, standards, and negative controls were run in technical duplicate.

### Gas chromatography

Gas chromatography measurements were used to determine the capacity of the three species to use sub-atmospheric concentrations of H_2_. Briefly, biological triplicate exponential phase or stationary phase cultures of each species were opened, equilibrated with ambient air (1 h), and resealed. These re-aerated vials were then amended with H_2_ (*via* 1% v/v H_2_ in N_2_ gas cylinder, 99.999% pure) to achieve final headspace concentrations of ~10 ppmv. Headspace mixing ratios were measured immediately after closure and at regular intervals thereafter until the limit of quantification of the gas chromatograph was reached (42 ppbv H_2_). For quantification, 2 mL headspace samples were measured using a pulsed discharge helium ionisation detector (model TGA-6791-W-4U-2, Valco Instruments Company Inc.) calibrated against ultra-pure H_2_ gas standards of known concentrations as described previously [18]. The vials for each species were maintained at their respective growth temperatures and agitation speeds for the entire incubation period to facilitate H_2_ and O_2_ transfer between the headspace and the culture. Concurrently, headspace mixing ratios from media-only negative controls (30 mL of media for each species) were measured to confirm that observed decreases in gas concentrations were biological in nature. First order rate constants (*k* values) for exponential and stationary phase H_2_ consumption were determined using the exponential function in GraphPad Prism (version 8.0.2).

### Phylogenetic analysis

A phylogenetic tree was constructed to investigate the distribution and evolutionary history of group 2a [NiFe]-hydrogenases across bacterial phyla. Amino acid sequences of the catalytic subunit of the group 2a [NiFe]-hydrogenase (HucL) and related enzymes were retrieved from the National Center for Biotechnology Information (NCBI) Reference Sequence (RefSeq) database by protein BLAST in February 2020. The resultant sequences were then classified using HydDB [26], with sequences matching group 2a [NiFe]-hydrogenases retained and any duplicate and multispecies sequences removed. The 207 amino acid sequences representative of genus-level diversity were aligned with reference sequences using Clustal W in MEGA X [47]. Evolutionary relationships were visualised by constructing a maximum-likelihood phylogenetic tree, with Neighbour-Join and BioNJ algorithms applied to a matrix of pairwise distances that were estimated using a JTT model and topology selected by superior log-likelihood value. Gaps were treated with partial deletion, the tree was bootstrapped with 500 replicates, and the tree was midpoint rooted. Sequences used in this analysis are listed in **Table S2**. Additionally, 20 annotated reference genomes (representative of order-level diversity) were retrieved from the NCBI GenBank database and manually analysed for putative group 2a [NiFe]-hydrogenase gene clusters. The web-based software Properon (doi.org/10.5281/zenodo.3519494) was used to generate to-scale gene organisation diagrams of these group 2a [NiFe]-hydrogenases.

## Results

### The expression profile of group 2a [NiFe]-hydrogenases is antithetical to group 1h [NiFe]-hydrogenases

We used qRT-PCR to quantify the expression of the large subunit of the group 2a [NiFe]-hydrogenase (*hucL*). The gene was expressed at moderate to high levels in all three strains during aerobic growth on preferred energy sources (organic carbon for *G. aurantiaca* and *C. aggregans*, ferrous iron for *A. ferrooxidans*) **(Fig. 1)**. Expression levels did not significantly differ between strains grown in an ambient air headspace containing atmospheric H_2_ or supplemented with 10% H_2_ **(Fig. 1)**. This suggests hydrogenase expression is constitutive and occurs even when atmospheric concentrations of the substrate are available.

**Figure 1.**
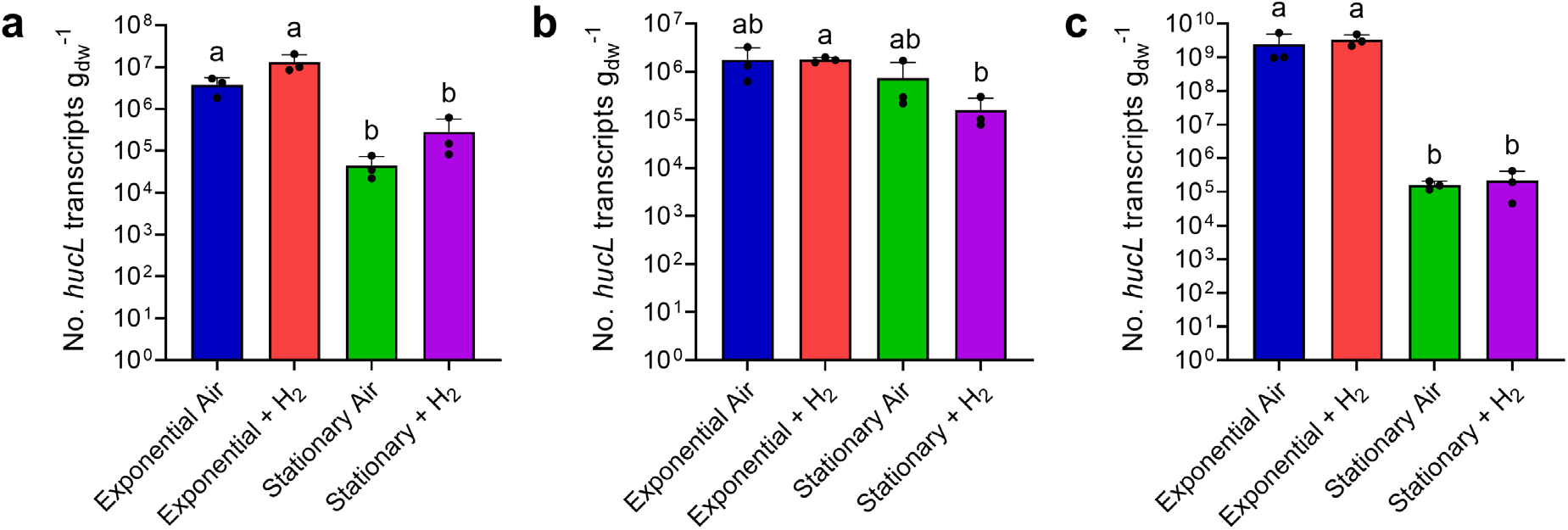
Expression of the group 2a [NiFe]-hydrogenase in three bacterial strains during growth and survival. The normalised transcript copy number of the large subunit gene (*hucL*) are plotted for **(a)** *Gemmatimonas aurantiaca* (locus GAU_0412), **(b)** *Acidithiobacillus ferrooxidans* (locus AFE_0702), and **(c)** *Chloroflexus aggregans* (locus CAGG_0471). Copy number was analysed by qRT-PCR in cultures harvested during exponential phase and stationary phase, in the presence of either ambient H_2_ or 10% H_2_. Error bars show standard deviations of three biological replicates (averaged from two technical duplicates) per condition. Values denoted by different letters were determined to be statistically significant based on a one-way ANOVA with post-hoc Tukey’s multiple comparison (*p* < 0.05).

Across all three strains, hydrogenase expression significantly decreased during the transition from growth to persistence. For *G. aurantiaca*, high expression was observed during exponential phase under both H_2_-supplemented and H_2_-unamended conditions (av. 8.4 × 10^6^ copies per g_dw_) and decreased 51-fold during stationary phase (av. 1.6 × 10^5^ copies g_dw_^−1^; *p* = 0.012) **(Fig. 1a)**. Hydrogenase expression of *A. ferrooxidans* was moderate during growth (av. 1.8 × 10^6^ copies per g_dw_) and dropped 3.9-fold in stationary phase cultures (av. 4.5 × 10^5^ copies per g_dw_; *p* = 0.013) **(Fig. 1b)**, whereas expression in *C. aggregans* was very high during exponential growth (av. 2.9 × 10^9^ copies g_dw_^−1^) and fell 15,000-fold during persistence (av. 1.9 × 10^5^ copies g_dw_^−1^; 0.003) **(Fig. 1c)**. Overall, while expression levels greatly vary between species, these results clearly show the group 2a [NiFe]-hydrogenase is expressed primarily in growing cells. These expression profiles contrast with the group 1h [NiFe]-hydrogenase, which is induced during long-term persistence in a range of species [10, 18, 20–23].

### Group 2a [NiFe]-hydrogenases oxidise H_2_ to sub-atmospheric levels

Hydrogenase activity of the three strains was inferred from monitoring changes in headspace H_2_ mixing ratios over time by gas chromatography. In line with the expression profiles **(Fig. 1)**, we observed that all three strains oxidised atmospheric H_2_ during growth in an ambient air headspace **(Fig. S1)**. These observations extend the trait of trace gas scavenging to three more species and suggest that group 2a [NiFe]-hydrogenases broadly have the capacity to oxidise H_2_ at atmospheric levels.

We subsequently monitored the consumption of H_2_ by exponential and stationary phase cultures in ambient air supplemented with 10 ppmv H_2_. For *G. aurantiaca* and *A. ferrooxidans*, H_2_ was oxidised to sub-atmospheric levels under both conditions in an apparent first-order kinetic process **(Fig. 2a** & **2b)**. However, biomass-normalised first-order rate constants were higher in exponential than stationary phase cells by 23-fold (*p* = 0.0029) and 120-fold (*p* < 0.0001) respectively **(Fig. 2d)**. For *C. aggregans*, H_2_ was oxidised at rapid rates in exponentially growing cells, but occurred at extremely slow rates in stationary cells **(Fig. 2c** & **2d)**. These observations support the qRT-PCR results by showing hydrogenase activity predominantly occurs during growth. It should be noted that additional [NiFe]-hydrogenases are encoded by both *C. aggregans* (group 3d) and *A. ferrooxidans* (group 1e and 3b). The additional hydrogenases are expressed at tenfold lower levels for *C. aggregans*, but at similar levels for *A. ferrooxidans*, and hence may contribute to H_2_ uptake **(Fig. S2)**. It is nevertheless likely that the group 2a [NiFe]-hydrogenases mediate atmospheric H_2_ uptake given (i) the H_2_ uptake activities of *C. aggregans* and *A. ferrooxidans* mimic that of *G. aurantiaca*, which lacks additional hydrogenases; (ii) previous genetic studies show group 2a enzymes mediate high-affinity aerobic H_2_ uptake in mycobacteria [12, 23]; and (iii) group 1e and 3b/3d enzymes are likely incapable of atmospheric H_2_ oxidation given their respective characterised roles in anaerobic respiration and fermentation [26].

**Figure 2.**
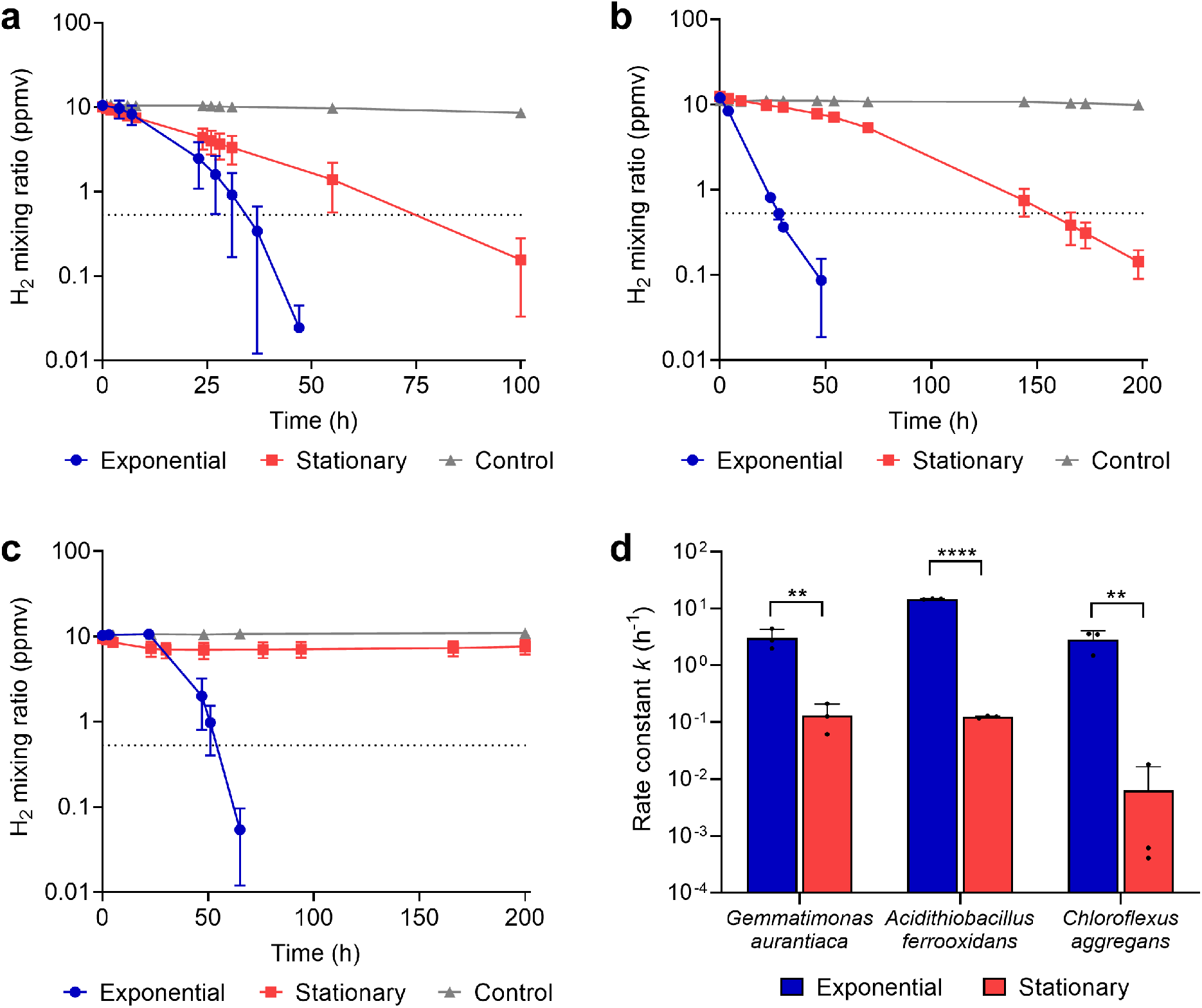
Hydrogenase activity in three bacterial strains during growth and survival. H_2_ oxidation by cultures of **(a)** *Gemmatimonas aurantiaca*, **(b)** *Acidithiobacillus ferrooxidans*, and **(c)** *Chloroflexus aggregans*. Error bars show the standard deviation of three biological replicates, with media-only vials monitored as negative controls. Dotted lines show the atmospheric concentration of hydrogen (0.53 ppmv). **(d)** Biomass-normalised first-order rate constants based on H_2_ oxidation observed in exponential and stationary phase cultures. Error bars show standard deviations of three biological replicates and statistical significance was tested using a two-way ANOVA with post-hoc Tukey’s multiple comparison (** = *p* < 0.01; **** = *p* < 0.0001).

### H_2_ consumption enhances mixotrophic growth in carbon-fixing strains

The observation that expression and activity of the group 2a [NiFe]-hydrogenase is optimal during growth suggests this enzyme supports mixotrophic growth. To test this, we monitored growth by optical density of the three strains in headspaces containing H_2_ at either ambient, 1%, or 10% mixing ratios. No growth differences in the obligate heterotroph *G. aurantiaca* were observed between the conditions (*p* = 0.30) **(Fig. 3a)**. In contrast, H_2_-dependent growth stimulation was observed for the obligate autotroph *A. ferrooxidans* (1.4-fold increase; *p* = 0.0003) **(Fig. 3b)** and facultative autotroph *C. aurantiaca* (1.2-fold increase; *p* = 0.029) **(Fig. 3c)**. This suggests that reductant derived from H_2_ oxidation can be used by these bacteria to fix CO_2_ through the Calvin-Benson and 3-hydroxypropionate cycles, respectively.

**Figure 3.**
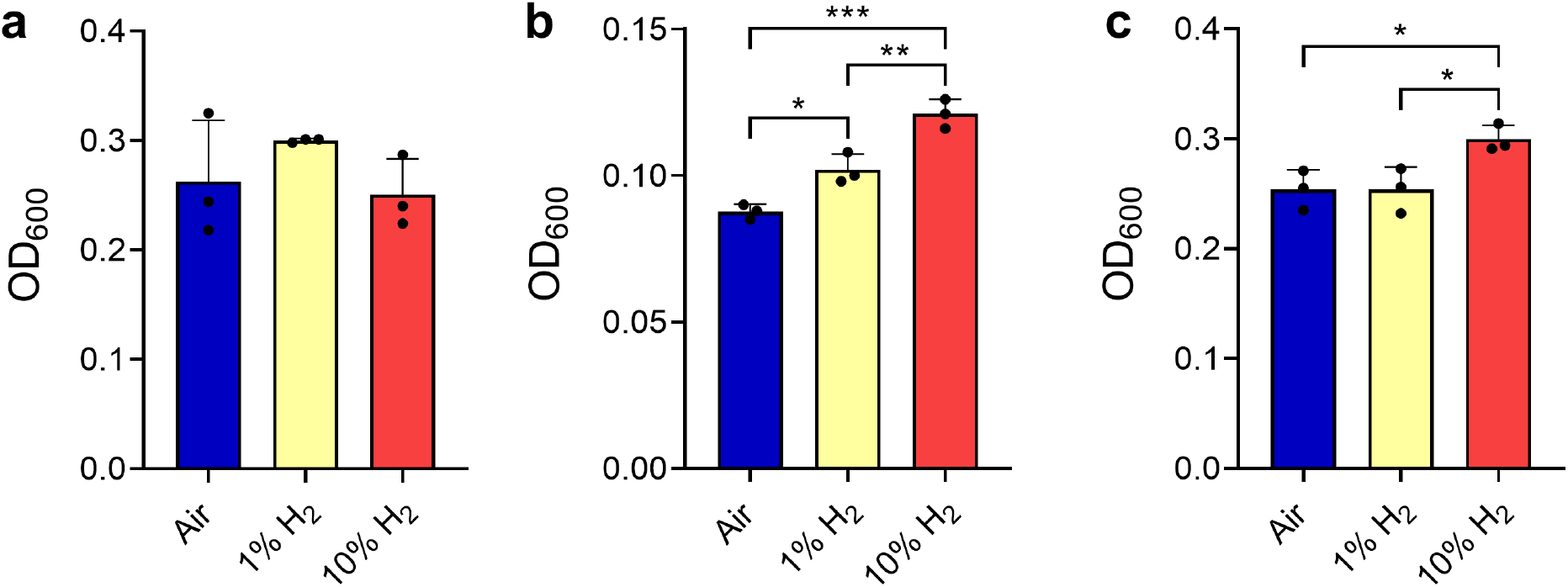
Effects of H_2_ supplementation on growth of three bacterial strains. The final growth yield (OD_600_) of **(a)** *Gemmatimonas aurantiaca*, **(b)** *Acidithiobacillus ferrooxidans*, and **(c)** *Chloroflexus aggregans* is shown in ambient air vials containing H_2_ at either ambient, 1%, or 10% concentrations. Error bars show the standard deviation of three biological replicates and statistical significance was tested using a one-way ANOVA with post-hoc Tukey’s multiple comparison (* = *p* < 0.05; ** = *p* < 0.01; *** = *p* < 0.001).

### Hydrogenases with common phylogeny and genetic organisation are widely distributed across 13 bacterial phyla

Finally, we surveyed the distribution of group 2a [NiFe]-hydrogenases to infer which other bacteria may oxidise atmospheric H_2_. We detected the large subunit of this hydrogenase (HucL) across 171 genera and 13 phyla **(Table S2**; **Fig. S3)**; this constitutes a 3.2-fold increase in the number of genera and 1.4-fold increase in the number of phyla reported to encode this enzyme [4, 26]. The HucL-encoding bacteria include various known hydrogenotrophic aerobes, such as *Nitrospira moscoviensis* (Nitrospirota) [31], *Hydrogenobacter thermophilus* (Aquificota) [48], *Kyrpidia tusciae* (Firmicutes) [49], *Sulfobacillus acidophilus* (Firmicutes) [50], and *Pseudonocardia dioxanivorans* (Actinobacteriota) [51], suggesting these strains may also consume atmospheric H_2_. The hydrogenase was also distributed in various lineages of Bacteroidota, Alphaproteobacteria, Gammaproteobacteria, and Deinococcota for which H_2_ oxidation has not, to our knowledge, been reported.

A maximum-likelihood phylogenetic tree showed the retrieved HucL sequences form a well-supported monophyletic clade. Most sequences clustered into four major radiations, Bacteroidota-associated, Cyanobacteria-associated, Proteobacteria-associated (including *A. ferrooxidans*), and a mixed clade containing sequences from seven phyla (including *G. aurantiaca* and *C. aggregans*) **(Fig. 4)**. Several genes were commonly genomically associated with *hucL* genes in putative operons, including the hydrogenase small subunit (*hucS*), a Rieske-type iron-sulfur protein (*hucE*) [34], hypothetical proteins (including NHL-repeat proteins) [33], and various maturation factors **(Fig. S4)**. The group 2a [NiFe]-hydrogenases are distinct in both phylogeny and genetic organisation to the two most closely related hydrogenase subgroups, the previously described group 2e [NiFe]-hydrogenases of aerobic hydrogenotrophic Crenarchaeota [26, 52] and the novel group 2f [NiFe]-hydrogenases that are distributed sporadically in bacteria and archaea **(Fig. 4)**.

**Figure 4.**
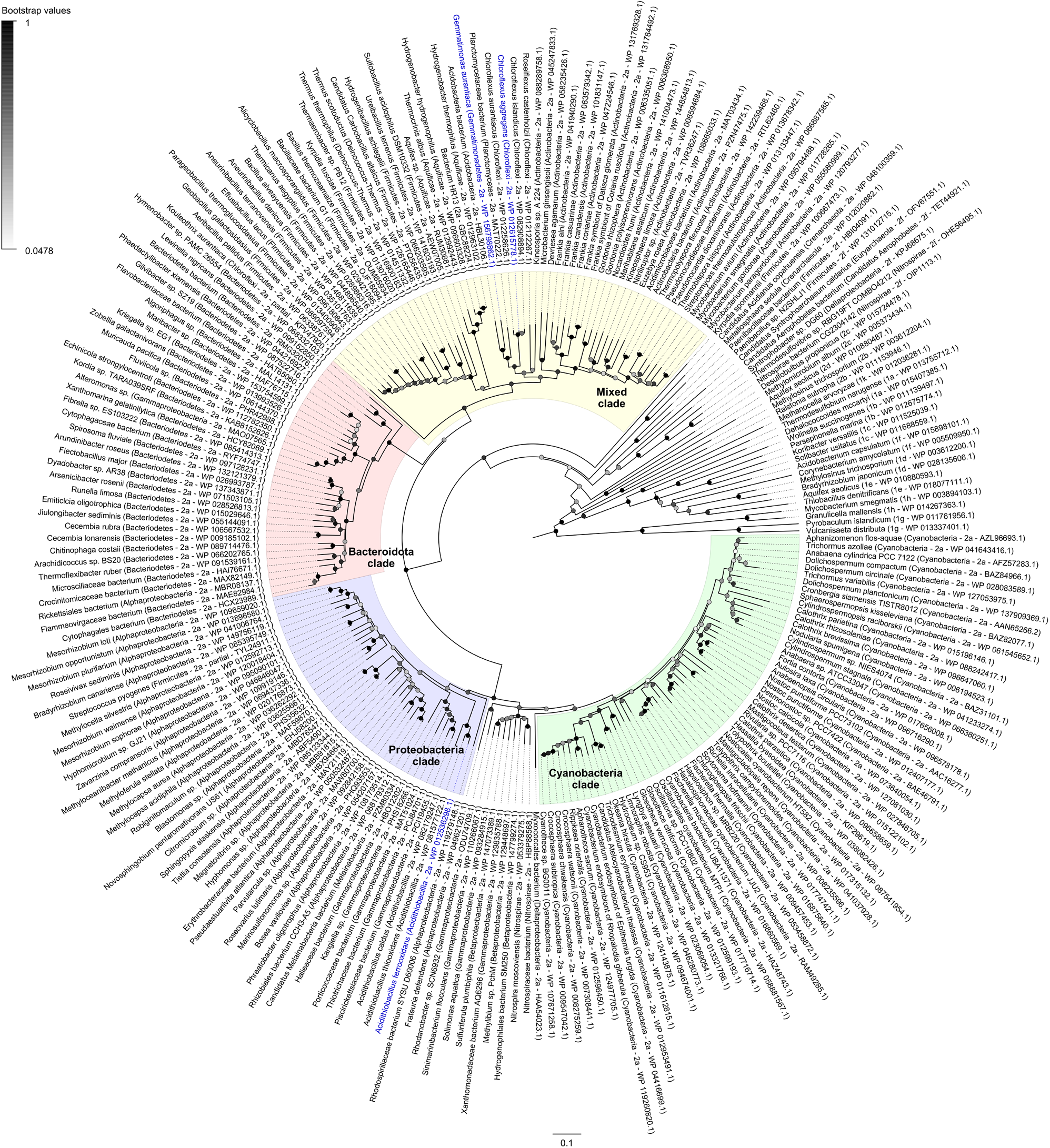
Radial phylogenetic tree showing the distribution and evolutionary history of the group 2a [NiFe]-hydrogenase. Amino acid sequences of the catalytic subunit of the group 2a [NiFe]-hydrogenase (*hucL*) are shown for 171 bacterial genera. The taxon names of the three study species, *G. aurantiaca*, *A. ferrooxidans*, and *C. aggregans*, are coloured in blue. The tree was constructed using the maximum-likelihood method (gaps treated with partial deletion), bootstrapped with 500 replicates, and rooted at the mid-point.

## Discussion

Overall, these findings overturn the paradigm that atmospheric H_2_ oxidation is primarily a persistence-linked trait. We infer that group 2a [NiFe]-hydrogenases are optimally expressed and active during exponential phase, consume H_2_ at sub-atmospheric concentrations, and support mixotrophic growth. Largely concordant findings were made in three phylogenetically, physiologically, and ecologically distinct bacterial species. These findings contrast with multiple pure culture studies that have linked expression, activity, and phenotypes associated with group 1h [NiFe]-hydrogenases to survival rather than growth [10, 12, 18, 20, 22, 24, 25]. However, a growth-supporting role of atmospheric H_2_ oxidation is nevertheless consistent with several surprising recent reports: the measurement of atmospheric H_2_ oxidation during growth of several strains [12, 19, 24, 53]; the discovery of an Antarctic desert community driven by trace gas oxidation [9]; and the isolation of a proteobacterial methanotroph thought to grow on air alone [54]. Together, these findings suggest that the current persistence-centric model of atmospheric H_2_ utilisation is overly generalised and that this process also supports growth.

Atmospheric H_2_ oxidation during growth is likely to primarily benefit bacteria that adopt a mixotrophic lifestyle. While atmospheric H_2_ alone can sustain bacterial maintenance, theoretical modelling suggests this energy source is insufficiently concentrated to permit growth as the sole energy source [1, 55]. Instead, bacteria that co-oxidise this dependable gas with other organic or inorganic energy sources may have significant selective advantages, especially in environments where resource availability is very low or variable. Likewise, it is probable that many bacteria in natural environments supplement growth by taking advantage of transient increases in H_2_ availability. For example, the metabolic generalist *C. aggregans* may facilitate its expansion in geothermal mats by simultaneously utilising geothermal and atmospheric sources of H_2_, in addition to sunlight and organic compounds [39, 40, 56]. Similarly, in the dynamic environment of wastewater treatment plants, *G. aurantiaca* may be well-suited to take advantage of fermentatively-produced H_2_ released during transitions between oxic and anoxic states [36, 57].

The ability to consume atmospheric H_2_ may also be particularly advantageous during early stages of ecological succession. Indeed, *A. ferrooxidans* may initially rely on this atmospheric energy source as it colonises barren tailings and establishes an acidic microenvironment conducive for iron oxidation [58]. Hydrogen synthesis in tailings can further benefit *A. ferrooxidans* as acid conditions and more complex bacterial consortia develop. Specifically, acetate-dependent growth of dissimilatory sulfate reducing bacteria in tailings [59] will initiate endogenous geochemical production of trace hydrogen (FeS + H_2_S → FeS_2_ + H_2_). As tailings cycle between aerobic (vadose) and anaerobic (water-saturating) conditions, the H_2_ available from atmospheric and geochemical sources respectively may provide a continuous energy source for *A. ferrooxidans*. In addition, any environments possessing sulfate and iron, i.e., ‘downstream’ from acid-generating ecosystems (including marine sediments), can generate hydrogen via bacterial sulfate reduction.

This study also identifies key microbial and enzymatic players in the global hydrogen cycle. The group 2a [NiFe]-hydrogenase is the second hydrogenase lineage shown to have a role in atmospheric H_2_ oxidation across multiple bacterial phyla. The group 1h enzyme is probably the main sink of the H_2_ cycle given it is the predominant hydrogenase in most soils [4, 11, 60]. However, the group 2a enzyme is moderately to highly abundant in many soil, marine, and geothermal environments [60], among others, and hence is also likely to be a key regulator of H_2_ fluxes. This study also reports atmospheric H_2_ oxidation for the first time in two globally dominant phyla, Proteobacteria and Gemmatimonadota, and uncovers *A. ferrooxidans* as the first H_2_-scavenging autotroph. Until recently, atmospheric H_2_ oxidation was thought to be primarily mediated by heterotrophic Actinobacteriota [1, 10–12], but it is increasingly apparent that multiple aerobic lineages are responsible [4, 17–19, 22, 34]. Some six phyla have now been described that are capable of atmospheric H_2_ oxidation and, given the group 2a [NiFe]-hydrogenase is encoded by at least eight other phyla, others will likely soon be described. It is possible that atmospheric H_2_ oxidation extends to other important groups, such as nitrite-oxidising Nitrospirota [31], methane-oxidising Proteobacteria [54], and potentially even oxygenic phototrophs; while Cyanobacteria are known to recycle endogenously-produced H_2_ [27, 62, 63], it should be tested whether they can also scavenge exogenous H_2_. Indeed, while atmospheric H_2_ oxidisers were only recently discovered [10, 14, 64], it is now plausible that these bacteria may represent the rule rather than the exception among aerobic H_2_ oxidisers.

## Footnotes

## Acknowledgements

This work was supported by an ARC DECRA Fellowship (DE170100310; awarded to C.G.), an ARC Discovery Grant (DP200103074; awarded to C.G. and R.G.), an NHMRC EL2 Fellowship (APP1178715; salary for C.G.), and Australian Government Research Training Program Stipend Scholarships (awarded to Z.F.I. and K.B.).

## Author contributions

C.G. and Z.F.I. conceived this study. C.G., Z.F.I., and R.G. supervised this study. C.G., Z.F.I., and C.W. designed experiments. Z.F.I., C.W., and K.B. performed experiments. Z.F.I., C.W., and C.G. analysed data. E.J.G. and G.S. contributed to study conception and experimental development. Z.F.I., C.G., and C.W. wrote the paper with input from all authors.

