## Supplementary information for "A widely distributed hydrogenase oxidises atmospheric H_2_ during bacterial growth"

### Table S1. Oligonucleotide primers used in this study.

| Primer | Sequence (5’ to 3’) | Tm (°C) |
|---|---|---|
| *G. aurantiaca* 16S_fwd (515F) | GTG YCA GCM GCC GCG GTA A | 54 |
| *G. aurantiaca* 16S_rvs (806rB) | GGA CTA CNV GGG TWT CTA AT | 54 |
| *G. aurantiaca* HucL_fwd | TGC ATG GAC CGA AGC AAG | 62 |
| *G. aurantiaca* HucL_rvs | AAT GAG CGT GGC GTT GTG | 62 |
| *C. aggregans* 16S_fwd | CGA AAG AAC CTT ACC CGG GC | 61 |
| *C. aggregans* 16S_rvs | CGA TCT GCA CTG AGA CCA CG | 61 |
| *C. aggregans* HucL_fwd | CAT CGA GGG GAG AAA TGC GG | 61 |
| *C. aggregans* HucL_rvs | GGA AGA GCG GGT CGT AGT CT | 61 |
| *C. aggregans* HoxH_fwd | CGA TCC GAT TGA CTA CCG CG | 61 |
| *C. aggregans* HoxH_rvs | GGC GCG AAG GTT AGG TGA AA | 61 |
| *A. ferrooxidans* rpoC_fwd | GGT GCA GCA GGA TTC GTT CA | 61 |
| *A. ferrooxidans* rpoC_rvs | ACG AGG GAG GTC ATG GGA AG | 61 |
| *A. ferrooxidans* HucL_fwd | GGT TGG GCA AGT ACG TCT CG | 61 |
| *A. ferrooxidans* HucL_rvs | GCG ATC AGT TGC CGG GAT AG | 61 |
| *A. ferrooxidans* HyiB_fwd | GGA GAG CAA GAT CAT CGC CG | 61 |
| *A. ferrooxidans* HyiB_rvs | GGG AGT TCG GTG CCT TTG AG | 61 |
| *A. ferrooxidans* HyhL_fwd | CCA AAG CCG TGA GAT CAG CC | 61 |
| *A. ferrooxidans* HyhL_rvs | TGT ATC CAC TCG CCG CAG TA | 61 |

### Table S2 (xlsx): Microbial distribution and amino acid sequences of the group 2a [NiFe]-hydrogenase large subunit HucL.

**Figure S1. Consumption of atmospheric $H_2$ during exponential growth of the three cultures.** $H_2$ mixing ratios of cultures of **(a)** *Gemmatimonas aurantiaca*, **(b)** *Acidithiobacillus ferrooxidans*, and **(c)** *Chloroflexus aggregans* are shown. Ratios were measured upon inoculation (blue bars), during mid-exponential growth (yellow bars), and in late stationary phase (red bars); the different sampling times between the cultures reflect their distinct growth parameters. Error bars show standard deviations of three biological replicates and statistical significance was tested using a two-way ANOVA with post-hoc Tukey's multiple comparison (** = $p < 0.01$; *** = $p < 0.001$; **** = $p < 0.0001$).

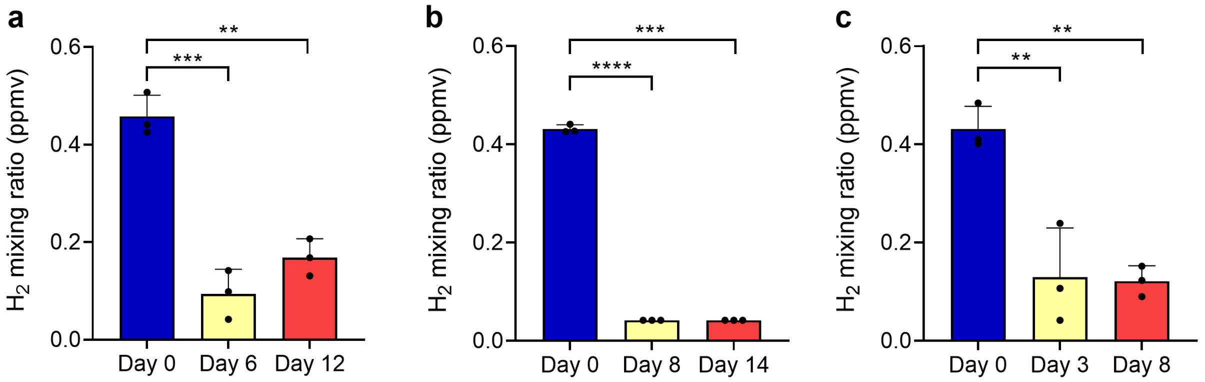

**Figure S3. Distribution of group 2a [NiFe]-hydrogenases across microbial genomes.** Results are based on the number of genomes in which the large subunit gene was detected (*hucL*) and are shown by phylum.

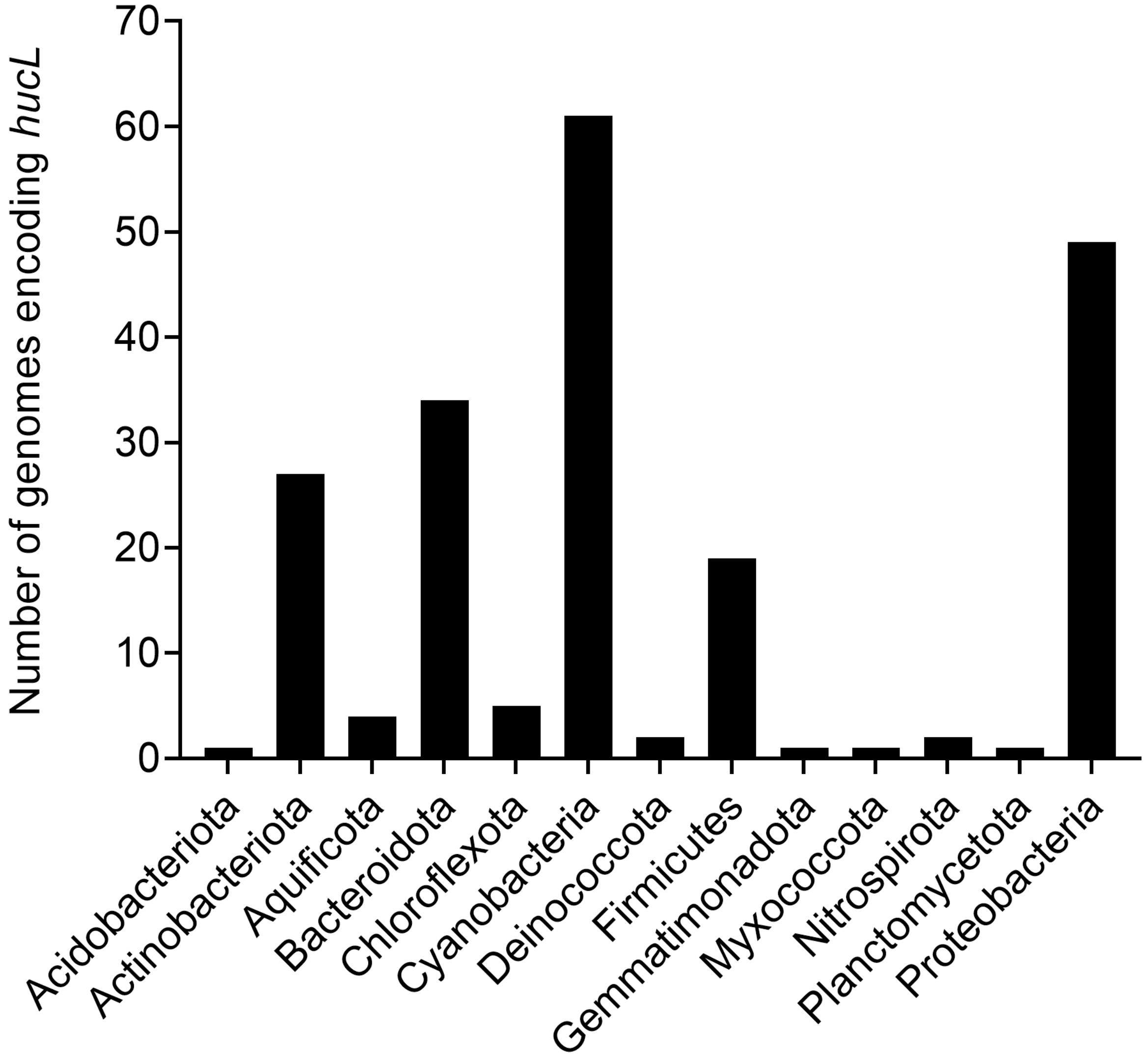

**Figure S4. Genetic organisation of group 2a [NiFe]-hydrogenases across ten phyla.** Abbreviations: HucL = hydrogenase large subunit; HucS = hydrogenase small subunit; HypABCDEF = hydrogenase maturation factors; HMP = hydrogenase maturation peptidase; NHL = NHL repeat protein; HP = conserved hypothetical protein; GmhA = putative phosphoheptose isomerase; Rieske = Rieske-like iron-sulfur protein (HucE); FAD = putative FAD-dependent oxidoreductase. Gene length is shown to scale and gene identifiers are as per the nomenclature of HydDB.

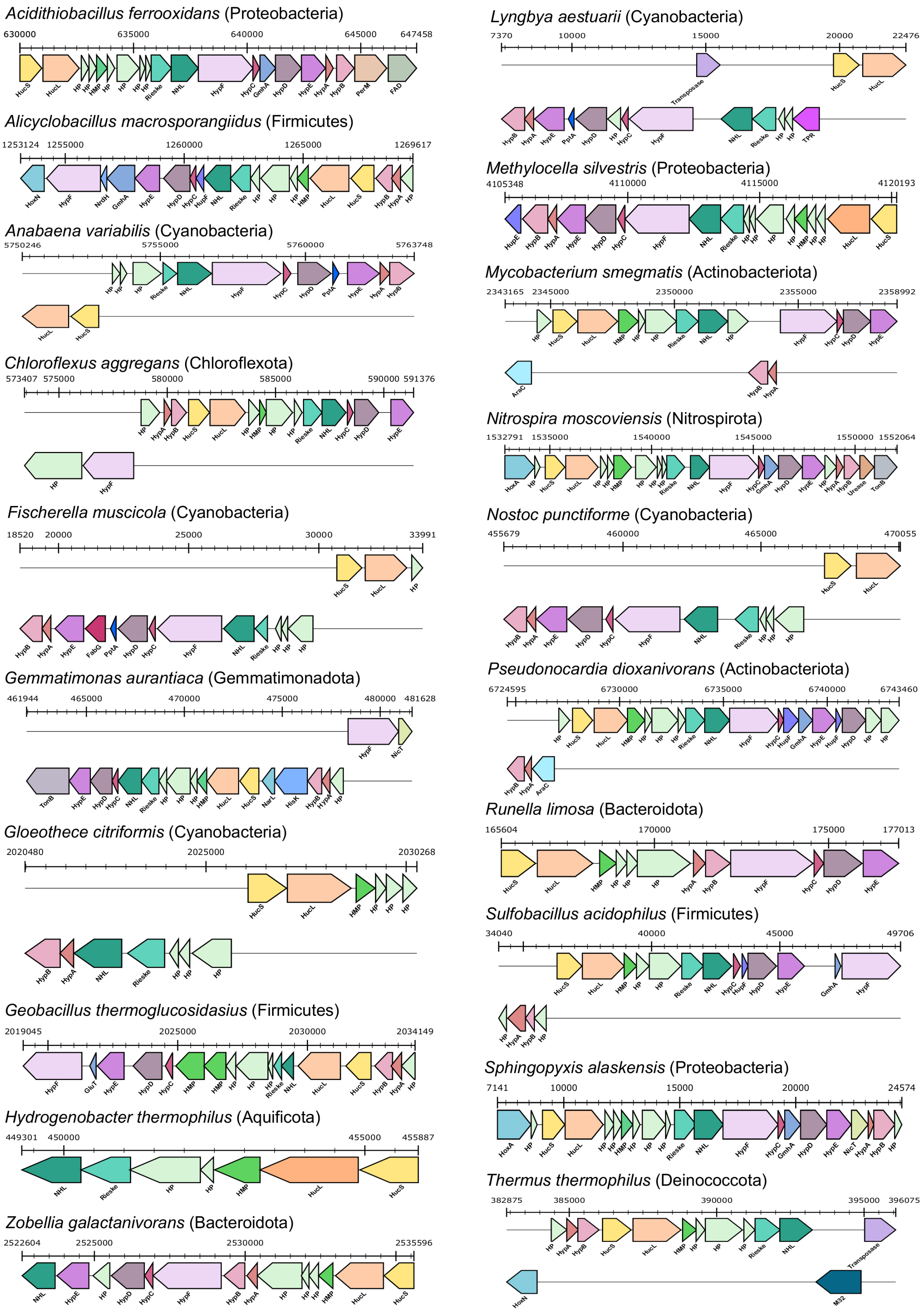

Acidithiobacillus ferrooxidans (Proteobacteria)
630000 635000 640000 645000 647458
Alicyclobacillus macrosporangiidus (Firmicutes)
1253124 1255000 1260000 1265000 1269617
Anabaena variabilis (Cyanobacteria)
5750246 5755000 5760000 5763748
Chloroflexus aggregans (Chloroflexota)
573407 575000 580000 585000 590000 591376
Fischerella muscicola (Cyanobacteria)
18520 20000 25000 30000 33991
Gemmatimonas aurantiaca (Gemmatimonadota)
461944 465000 470000 475000 480000 481628
Gloeothece citriformis (Cyanobacteria)
2020480 2025000 2030268
Geobacillus thermoglucosidasius (Firmicutes)
2019045 2025000 2030000 2034149
Hydrogenobacter thermophilus (Aquificota)
449301 450000 455000 455887
Zobellia galactanivorans (Bacteroidota)
2522604 2525000 2530000 2535596
Lyngbya aestuarii (Cyanobacteria)
7370 10000 15000 20000 22476
Methylocella silvestris (Proteobacteria)
4105348 4110000 4115000 4120193
Mycobacterium smegmatis (Actinobacteriota)
2343165 2345000 2350000 2355000 2358992
Nitrospira moscoviensis (Nitrospirota)
1532791 1535000 1540000 1545000 1550000 1552064
Nostoc punctiforme (Cyanobacteria)
455679 460000 465000 470055
Pseudonocardia dioxanivorans (Actinobacteriota)
6724595 6730000 6735000 6740000 6743460
Runella limosa (Bacteroidota)
165604 170000 175000 177013
Sulfobacillus acidophilus (Firmicutes)
34040 40000 45000 49706
Sphingopyxis alaskensis (Proteobacteria)
7141 10000 15000 20000 24574
Thermus thermophilus (Deinococcota)
382875 385000 390000 395000 396075
